# Stem cell derived human microglia transplanted in mouse brain to study genetic risk of Alzheimer’s Disease

**DOI:** 10.1101/562561

**Authors:** Renzo Mancuso, Johanna Van Den Daele, Nicola Fattorelli, Leen Wolfs, Sriram Balusu, Oliver Burton, Annerieke Sierksma, Yannick Fourne, Suresh Poovathingal, Amaia Arranz-Mendiguren, Carlo Sala Frigerio, Christel Claes, Lutgarde Serneels, Tom Theys, V. Hugh Perry, Catherine Verfaillie, Mark Fiers, Bart De Strooper

## Abstract

Genetics highlight the central role of microglia in Alzheimer’s disease but at least 36% of AD-risk genes lack good mouse orthologues. Here, we show that embryonic stem cell (ESC)-derived human microglia successfully engraft the mouse brain and recapitulate transcriptionally primary human microglia derived from human surgical samples. Upon exposure to oligomeric Aβ a wide range of AD-risk genes are expressed that are not readily studied in current mouse models for AD. This work provides a unique humanized animal model that will allow elucidating the role of genetic risk in the pathogenesis of AD.

## Main text

Genome-wide association studies (GWAS) have highlighted the central role of the microglia response in determining the risk of Alzheimer’s disease (AD)^1–6,7^. We constructed a list of 82 AD risk genes by combining 37 established GWAS genes^3,8^ and 45 genes with potential contribution to the pathology selected from Marioni et al.^4^ (significance threshold of p<5×10^−8^) (Supplementary Table 1). Strikingly, twenty of these genes do not have a clear mouse orthologue (Fig. 1a and Supplementary Table 1), including *CR1*, *CD33*, *APOC*, *MS4A4*, etc. In addition, 10 more AD risk genes are less than 60% similar between mouse and human at the primary amino acid sequence (Ensembl build 95^9^), including the major AD-risk gene *TREM2* (Fig. 1a and Supplementary Table 1). Another major difference is the *APOE* polymorphism which is the most frequent genetic risk factor for sporadic AD and does not exist in rodents. At gene expression level, while mouse and human microglial transcriptomes might appear conserved, mouse microglia display less expression of AD- and Parkinson Disease risk associated genes^10^, and the divergence between species is exacerbated with ageing^11^. This clearly shows that mouse microglia have limitations as a model to study AD genetic risk. Although it is possible to culture human microglia *in vitro*, it is very difficult to maintain them with a homeostatic signature^12^.

**Figure 1.**
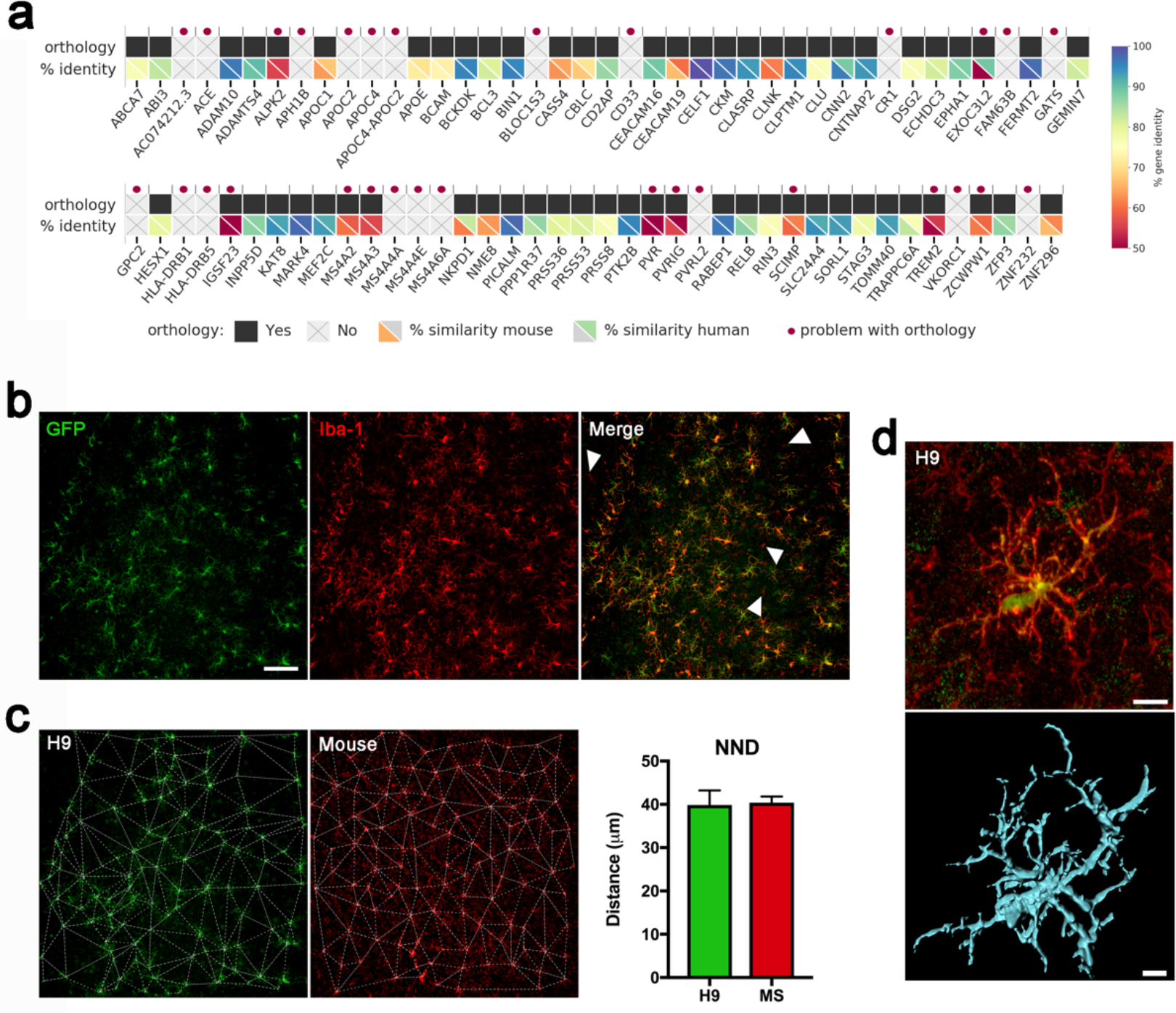
Human ESC-derived microglia successfully engraft the mouse brain. (**a**) Limited orthology between human and mouse AD-related genes. From a total of 82 AD risk genes including 37 established GWAS genes^3,8^ and other genes with potential contribution to the pathology (selected from Marioni et al.^4^, setting a threshold of p<5×10^−8^), 30 (marked with a red dot) do not have a clear mouse orthologue or display low percentage of similarity between mouse and human at the primary amino acid sequence (**b**) H9-microglia successfully engraft the mouse brain and (**c**) distribute across the parenchyma forming a mosaic with similar nearest neighbour distance to that of mouse cells from adjacent areas (n=4 per group, two-tailed t-test p>0.05, graph shows mean±SEM). H9-microglia are labelled in green (Iba1^+^ GFP^+^), whereas arrowheads highlight mouse cells (Iba1^+^ GFP^−^) co-existing with H9-microglia in the grafted areas of the parenchyma. Scale bar, 100μm (**d**) Higher magnification microphotographs and 3D reconstruction by Imaris show resting morphology with high complexity branching in H9-microglia. Scale bar 5μm.

Here, we investigate to what extent embryonic pluripotent stem cells (ePS) (H9-ESC)-derived human microglia transplanted into mouse brain mimic morphological and transcriptomic features of primary human microglia, and how mouse and human microglia respond differently to an AD related insult.

We first analysed survival and integration of H9-microglia in the mouse brain. H9-ESCs were differentiated into monocytes^13^, purified, and further differentiated into microglia using cytokines CSF1, IL-34, TGF-β and CX3CL1 (Supplementary Fig. 1). We transplanted H9-microglia into the brain of postnatal day 4 (P4) *Rag2*^−/−^ *Il2rγ*^−/−^ *hCSF1*^*KI*^ mice^14^, abbreviated to *hCSF1^KI^*. Human microglia require hCSF1 for their growth and survival^14^. We treated *hCSF1*^*KI*^ mice with the Colony-Stimulating Factor 1-Receptor (CSF1R) inhibitor BLZ945^15^, 24h and 48h prior to the transplantation to remove 50-60% of mouse microglia (Supplementary Fig. 2), providing the human cells a permissive environment for integration. Eight weeks after transplantation, H9-microglia showed a widespread distribution across multiple areas of the brain, including striatum, rostral hippocampus and multiple fibre tracts (Supplementary Fig. 3). H9-microglia showed a complex ramified morphology, and a mosaic distribution with a nearest neighbour distance (NND)^16^ of 40μm, similar to mouse cells (n=4 mice per group, Fig. 1b-d).

**Figure 2.**
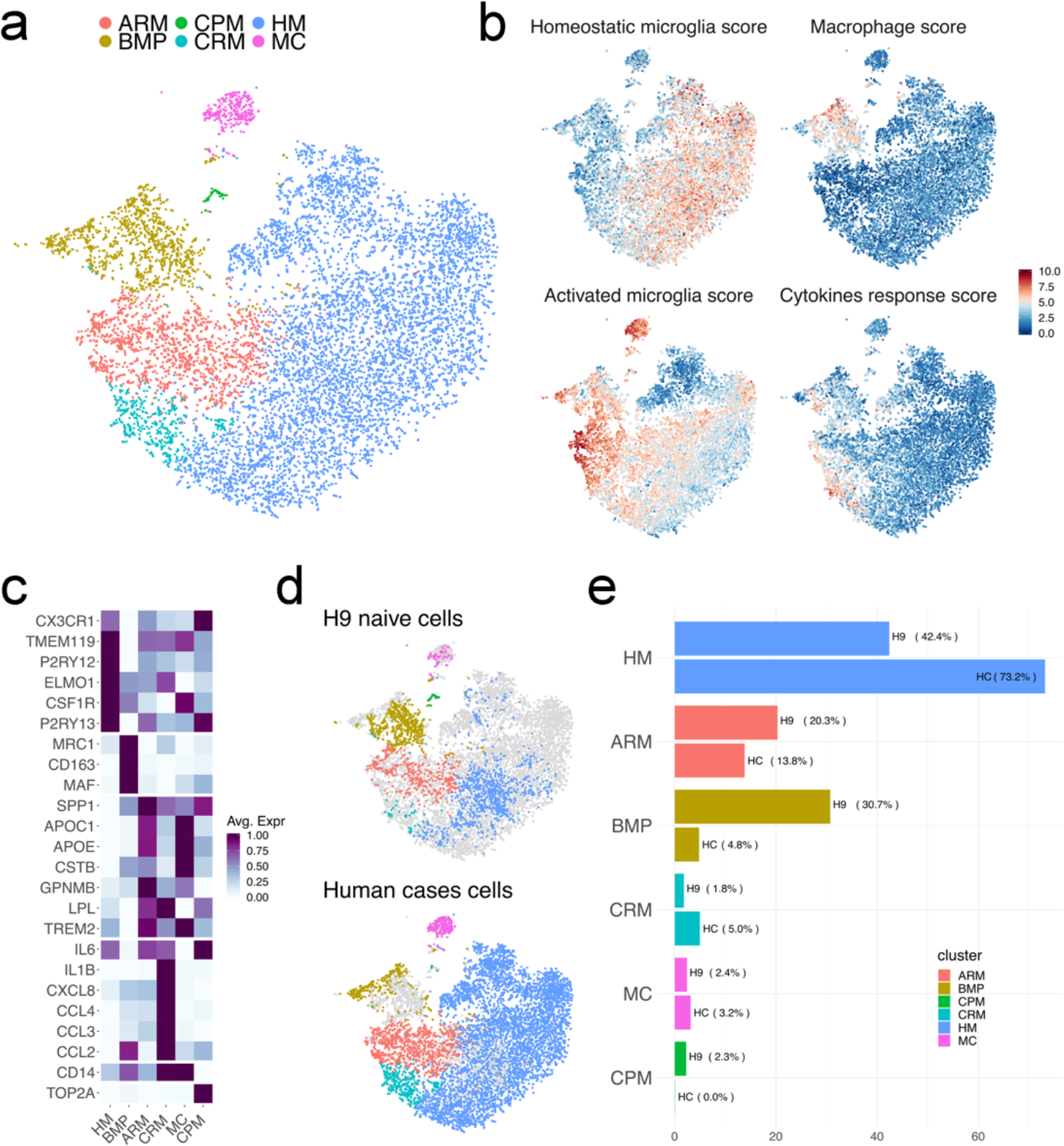
H9-microglia isolated 8 weeks after transplantation are similar to human primary microglia. (**a**) t-SNE plot visualizing 10,512 cells sorted based on CD11b (primary human), or CD11b hCD45 and GFP (H9-microglia) staining (2,469 H9- and 8,043 primary cells) after quality control, and removal of peripheral cells. Cells are coloured according to clusters identified with Seurat’s kNN. (**b)** t-SNE plots as in **a**, coloured by the combined level of expression of groups of genes that characterise distinct microglial states^17,19^ (and Sala-Frigerio, submitted, manuscript provided for the referees only) (see Supplementary table 3). (**c**) Expression of the most expressed genes from the different clusters observed in **a**. (**d**) t-SNE plot as in **a**, coloured by the distribution of H9 and primary microglia. (**e**) Percentage of cells from either H9 or primary human microglia across the different clusters identified.

**Figure 3.**
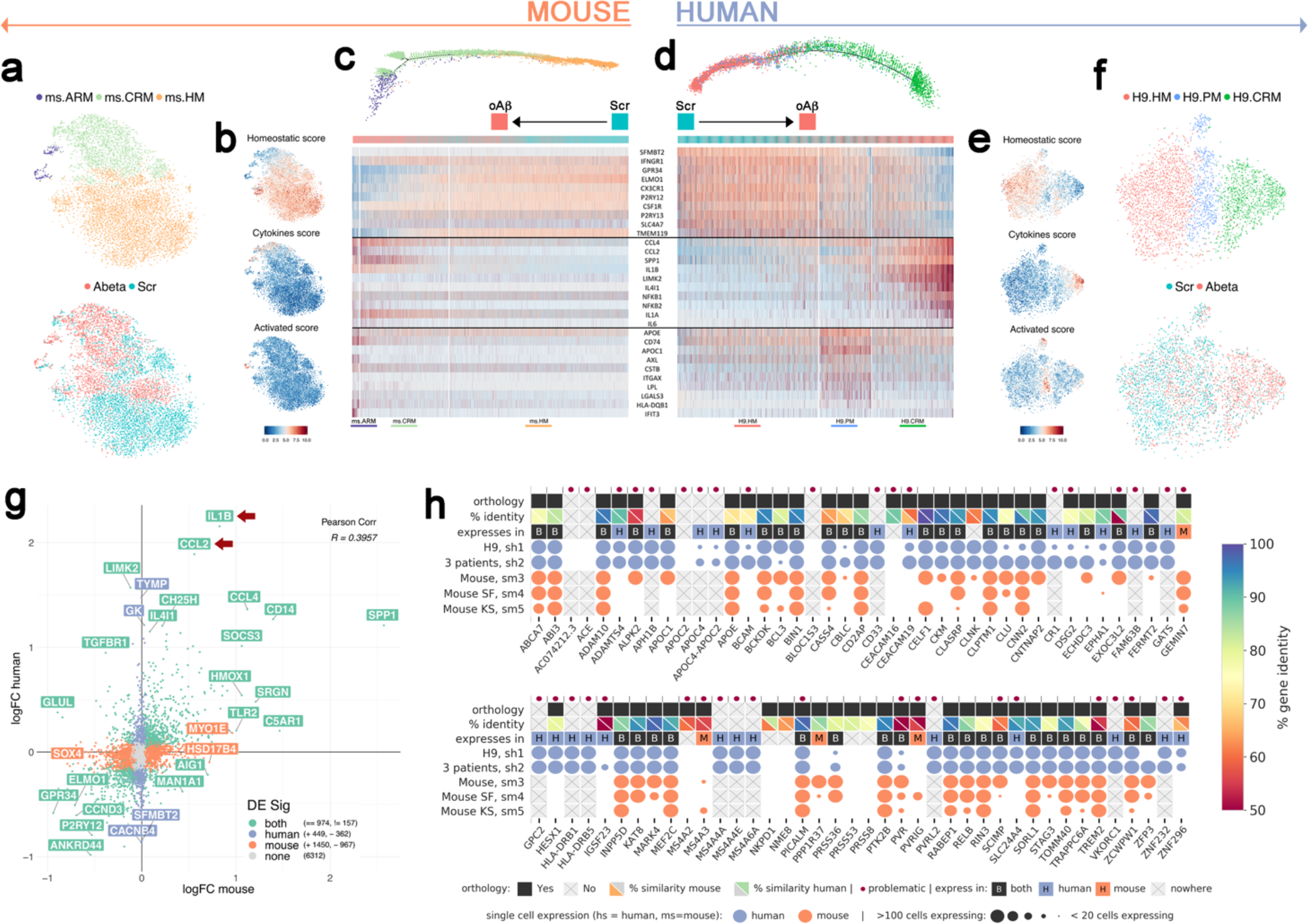
Human and host mouse microglial response to oligomeric Aβ. (**a-c**) Analysis of the response of endogenous (*Rag2*^−/−^) mouse microglia upon oligomeric Aβ challenge. (**a**) t-SNE plot visualizing the 9942 endogenous mouse microglia passing quality control, and after removal of peripheral cells, brain resident macrophages (BMP) cycling cells and doublets. Cells are coloured according to clusters identified with Seurat’s kNN (upper panel; ms.HM: (mouse) Homeostatic Microglia, ms.CRM: Cytokine Response Microglia, ms.ARM: Activated Response Microglia) and treatment (lower panel; Scr: scrambled peptide, oAb: oligomeric Ab). (**b**) t-SNE plots as in **a**, coloured by the combined level of expression of groups of genes that characterise distinct microglial states. Expression of the main single genes used to construct the different profiles is in Supplementary Fig. 8 and Supplementary table 3. (**c**) Plot of the phenotypic trajectory followed by endogenous mouse microglia upon oligomeric Ab exposure, obtained by an unbiased pseudotime ordering with Monocle 2 and coloured by clusters as in **a**. Mouse microglia followed a linear trajectory from ms.HM to ms.CRM to ms.ARM. The heatmap shows the differential expression of representative genes from each cluster, ordered by pseudotime. (**d-f**) Analysis of the response of H9-mouse microglia upon oAβ. (**f**) t-SNE plot visualizing the 4880 H9 microglia passing quality control, and after removal of brain resident macrophages, cycling cells and doublets. Cells are coloured according to clusters identified with Seurat’s kNN (upper panel; H9.HM: Homeostatic Microglia, H9. PM: Primed Microglia, H9.CRM: Cytokine Response Microglia) and treatment (lower panel; Scr: scrambled peptide, oAβ: oligomeric Ab). (**e)**t-SNE plots as in **a**, coloured by the combined level of expression of groups of genes that characterise distinct microglial states. Expression of the main genes used to construct the different profiles is in Supplementary Fig. 9). (**d**) Plot of the phenotypic trajectory followed by H9-microglia upon oligomeric Aβ exposure, obtained by an unbiased pseudotime ordering with Monocle 2 and coloured by clusters as in **f**. H9-microglia followed a linear trajectory from H9.HS and H9.PM, to H9.CRM. The heatmap shows the differential expression of representative genes from each cluster, ordered by pseudotime. (**g**) Correlation analysis of the log-fold change (logFC) in H9 (y-axis) and host (*Rag2*^−/−^) mouse (x-axis) microglia upon oligomeric Aβ challenge relative to scrambled peptide (Pearson correlation, R=0.3957). Differentially expressed genes are highlighted in green when significant in both species, blue only in H9 or orange only in mouse microglia. (**h**) Extension of the table shown in Fig. 1a highlighting the limited orthology between AD risk genes in humans and mice (30 genes highlighted by red dots). Expression profile of 82 AD liked genes in our datasets (H9-microglia, sh1; primary human microglia from 3 patients, sh2; and mouse host microglia, sm3), and wild type mouse microglia from 2 independent datasets of 12-week-old immunocompetent C57Bl/6 mice (Sala-Frigerio, submitted, manuscript provided for the referees only, SF, sm4; and Keren-Shaul et al., KS, sm5^17^). We identified 25 genes with observed expression in human (sh1 and sh2) but not mouse (sm3, sm4 and sm5) microglia and, from those, 23 were observed in H9-microglia (sh1). This highlights the importance of using human systems to underpin the contribution of some genetic risk components in AD.

We compared the *in vivo* single cell transcriptomic profile of H9-derived microglia with that of human primary microglia obtained from cortical surgical resections (Supplementary Table 2). We sequenced in total 10,512 single cells comprising 2,469 H9-derived microglia from the 8 weeks old engrafted mouse brains and 8,043 primary human cells obtained from 3 patients (Supplementary Fig. 4 and 5 and online Methods). Overall, the microglia of the three patients’ samples showed different but clearly overlapping expression profiles and the H9-derived microglia were intermingled (Fig. S5c). We identified three main distinct populations of microglia (MG), and in addition clusters of brain resident macrophages (BMP) and monocytes (MC) (Fig. 2a, b and Supplementary Fig. 5). Within the microglial cluster, we observed homeostatic response microglia (HM) defined by *CX3CR1*, *TMEM119*, *P2RY12*, *ELMO1* and *P2RY13*; activated response microglia (ARM) enriched in *APOE*, *SPP1*, *CSTB*, *LPL* and *TREM2*^17^(Sala-Frigerio, submitted, manuscript provided for the referees only); and cytokine response microglia (CRM) expressing cytokine/chemokines such as *IL6*, *IL1B*, *CCL2*, *CCL3*, *CCL4*, (Figure 2b, c and Supplementary Fig. 5)^17–21^. The homeostatic microglia cluster HM was composed of a combination of cells from all human samples, while the ARM and CRM clusters contained also cells from the different patients, but were enriched with cells from Human Case 1 (Fig. 2d and Supplementary Fig. 5). Tissue from Human Case 1 was collected from a second intervention (Supplementary Table 2), which might underlie this observation. Strikingly, the transcriptome of H9-microglia overlapped with either the HM (42%) or the ARM (20%) and CRM clusters (1.8%) (Fig. 2e), whereas the remaining H9 cells clustered together with brain-resident macrophages (31%). A very small proportion of H9 cells (2.4%) showed a transcriptome resembling monocytes. Thus, transplanted H9-microglia cells in mice acquire transcriptional identities that fully recapitulate those of primary human microglia. At this stage of research, it is not possible to interpret the significance of the changes in relative proportions of different microglia cell states in the patients and the H9-transplants.

Next, we challenged H9-microglia in the mouse brain with an acute AD-related challenge, and compared their reaction with that of the endogenous host murine microglia. It is clear that the lack of interaction with the adaptive immune system is a limitation of the model and may modify the response of both H9 and host mouse microglia^22^. The *Rag2*^−/−^ host background might also affect the mouse microglia in other ways, for instance because of (unknown) developmental abnormalities inherent to the genetic background. Previous work however has reported no changes in microglial number, cytokine levels, and gene expression profiles between wild type and *Rag2*^−/−^ mice^22^, and we also investigated different independent microglial datasets from 8-12 weeks old immunocompetent mice and did not detect expression of *Rag2*^17,23^. We therefore compared the mouse and human microglia response on an oligomeric Aβ challenge in this model.

We excluded brain resident macrophages and other immune or cycling cells (see Supplementary Fig. 6). Oligomeric Aβ 1-42 (oAβ, 5μl 10μM) or scrambled peptide (Scr, 5μl 10μM) were prepared as previously^24^ (Supplementary Fig. 7) and injected in the ventricle of transplanted mice at 8-10 weeks post transplantation. We previously showed induction of cognitive alterations 24 hours after injection.^25^ We pooled tissue from two mice for each experimental condition (see online Methods). We isolated and sequenced 9942 endogenous host mouse cells and 4880 transplanted H9-microglia 6 hours after the oAβ/Scr injection. The endogenous mouse cells displayed 3 distinct clusters which we called homeostatic (HM), cytokine (CRM) and activated (ARM) response cells. The HM cluster was characterised by expression of *Cx3cr1*, *Tmem119*, *P2ry12*, *Elmo1* and *P2ry13*, whereas the ARM and CRM clusters showed a downregulation of homeostatic markers and increased expression of inflammatory cytokines/chemokines as above. The ARM cluster showed resemblance to the cluster we previously described in *Rag2*^*wt/wt*^ mouse microglia (Sala-Frigerio, submitted, manuscript provided for the referees only) and partially overlaps with the disease-associated microglia or DAM response (Fig. 3a-c and Supplementary Fig. 8)^18–21^. The homeostatic HM cluster was enriched in cells from mice treated with scrambled peptide (70%), whereas the cytokine CRM and activated ARM clusters mostly consisted of cells from the mice treated with oAβ (69% and 77%, respectively) (Fig. 3a). To understand the progression of the phenotypical change elicited by oAβ in the endogenous mouse host microglia, we performed a trajectory analysis with Monocle 2^26–28^. This showed a clear linear progression from HS to CRM to ARM response cells (Fig. 3c and Supplementary Fig. 8), suggesting that the 2 different activation states are part of a single successive response to oAβ in the mouse host (*Rag2*^−/−^) microglia.

We performed the same analysis on the human H9-microglia (excluding the brain resident macrophages and cycling cells, see Supplementary Fig. 6). Clustering revealed 3 different populations of homeostatic (H9.HM), primed (H9.PM), and cytokine (H9.CRM) expressing cells (Fig. 3d-f). Despite showing high expression of *APOE*, the “primed” H9.PM cluster was very different in gene expression profile from the ARM response in the mice host *Rag2*^−/−^ microglia, but also from the ARM response as we previously identified in *Rag2*^*wt/wt*^ mouse microglia (Sala-Frigerio, submitted, manuscript provided for the referees only) and consisted of a higher proportion of cells coming from the scrambled (65%) vs. oAβ cells (36%). This cluster expresses mild levels of homeostatic genes, as well as genes associated to microglial activation (e.g. *APOE*, *APOC1*, *CD74*, etc.) (Fig. 3d-f and Supplementary Fig. 9)^17–21^. H9.PM seems to encompass the response of the microglia to the experimental manipulation, i.e. the injection of peptides into the mice. The other two clusters are more similar to the clusters already defined above. The homeostatic H9.HM cluster was enriched with cells treated with scrambled peptide (68%) and expressed *CX3CR1*, *TMEM119*, *P2RY12*, *ELMO1* and *P2RY13*, (Fig 3d-f and Supplementary Fig. 9). On the other hand, the cytokine cluster H9.CRM, was enriched with cells from the oAβ treated mice (75%), and expressed high levels of multiple inflammatory cytokines and chemokines, such as *IL1B*, *IL6*, *CCL2*, *CCL4*, etc. (Fig. 3d-f). Trajectory inference revealed that oAβ induces a gradual phenotypical change of H9-microglia, from homeostatic and towards the cytokine-response state (Fig. 3d). Interestingly, microglia from the H9.PM cluster were enriched in the initial and middle phases of the phenotypical progression, indicating that these cells may represent a primed or transitioning state towards activation (Fig. 3d and Supplementary Fig. 9).

Despite the caveats discussed above, we considered of interest to compare the transcriptional response displayed by mouse (*Rag2*^−/−^) host and human H9-microglia to oAβ in order to pinpoint potential differences that may have relevant implications for disease and settle the bases for future in depth investigation. Using Ensembl Biomart, we extracted 10,914 one-to-one, bidirectional orthologues between the two species (Supplementary Table 2)^29^, and performed a correlation analysis comparing the log-fold change (logFC) in response to oAβ. Mouse and human cells showed a low correlation in their response to oAβ (Pearson correlation, R=0.4). Further pathway enrichment analysis on the differentially expressed genes upon oAβ revealed that a substantial proportion of the human response was involved in immune pathways (Supplementary Fig. 10a). Although upregulated in both species, the human response was particularly strong for *IL1B* and *CCL2* (red arrows in Fig. 3g), which have been experimentally implicated in the pathology of AD, particularly the induction of tau phosphorylation in neurons^30,31^ and infiltration of circulating inflammatory cells^32^ (Fig. 3g and Supplementary Fig. 10). Remarkably, 207 genes changed in either mouse or human, or showed opposite behaviour between mouse and human microglia (Supplementary Fig. 10): 112 genes were specifically upregulated in H9 but not mouse microglia (with a threshold logFC of 0.2). Genes differentially expressed in H9-but not host mouse microglia in response to oAβ included *TYMP*, described in aged primed microglia^20^; classical inflammatory genes such as *NFKB2;* or *PPARG*, which regulates APOE expression (Supplementary Fig. 10c). Examples of genes showing opposite response in human and mouse cells are *LIMK2*, associated with phagocytosis and previously shown to be upregulated by oAβ in primary human microglia in vitro^33^; *BIN1* and *PICALM*, GWAS AD-associated genes^3,8^; or *TGFBR1* (Supplementary Fig. 10c). Interestingly, *TGFBR1* is considered a classical homeostatic microglia marker in mouse microglia^19^, but was significantly upregulated in H9-microglia upon oAβ, indicating a radical different role in the human context (Supplementary Fig. 10 b, c). Activation of these pathways in H9-microglia was also accompanied by a more pronounced downregulation of multiple classical homeostatic markers, including *CX3CR1* and *P2RY12* (Supplementary Fig. 10b). Taken together, these results show that human and mouse (*Rag2*^−/−^) microglia display significant differences in their response to oAβ.

To further investigate the difference between human and mouse microglia, and also controlling for possible effects from the genetic background of the host mice, we compared the expression profiles of the 82 AD risk genes selected above (Fig. 1a) in the human and mouse microglia datasets generated here and in 2 independent single cell microglia data sets of immunocompetent 12 weeks old C57Bl/6 mice generated previously (Fig. 3h and Supplementary Table 1)^17^(Sala-Frigerio, submitted, manuscript provided for the referees only). We observed that 25 of the 30 genes identified in Fig. 1a as problematic to study in mouse models for various reasons, are actually expressed in human microglia isolated from patients, and 23 of these (92%) were confirmed to be expressed in the transplanted human H9-microglia (such as *APOC*, *CD33*, *CR1*, *MS4A*, *PVRL2* or *TREM2*). This emphasizes the importance of using human specific systems to interrogate genotype-phenotype interactions of the GWAS identified AD risk genes (Fig. 3h). As shown in Fig. 3h, expression of AD risk genes in *Rag2*^−/−^ vastly overlaps with that of *Rag2*^*wt/wt*^ microglia form independent datasets, which further supports our claim above regarding the relevance of comparing the host and transplanted microglia in our mouse model.

Although *in vitro* studies may provide some mechanistic insights into the function of human microglia, it is also clear that signals from the CNS microenvironment are required to sustain microglial specification^12^, and that a loss of those cues dramatically disrupts the microglia phenotype^19,10^. In addition, some AD-linked genes (e.g TREM2-membrane phospholipids/APOE, CD33-sialic acid, etc.) play a role in the cross-talk between microglia and other brain cells. The main challenge is to understand this cellular phase of AD^34^ and therefore introducing those complex aspects into a model of disease is extremely important. We present here a novel model of ESC-derived human microglia transplantation into the mouse brain that represents a step forward from previous efforts into that direction^35,36^, providing the human cells with the crucial environment that defines microglial identity. Given the limited similarity between mouse and human microglia in terms of AD-associated genes as discussed here, this model provides a very useful alternative to study the response of human microglia *in vivo* in the context of AD, opening important new routes to understand the role of the many genes identified in the GWAS and other genetic studies^3,8^ which are not well modelled in mouse cells as demonstrated here.

We are aware that the current model provides only proof of concept with ePS derived microglial cells, and that it is important to expand this model to the usage of patient derived induced Pluripotent Stem Cells (iPSC). Our preliminary data show however that this might come with additional hurdles, as not all cell lines (e.g. BJ1-iPSC, unpublished results) engraft equally successfully into the mouse brain. Further work is needed to explore the critical parameters and careful comparison of multiple iPS cell lines with the ePS line used here as benchmark. Regardless, the novel mouse chimeric model that we present here in combination with Crisper/Cas9 technology, opens unanticipated possibilities to model human specific aspects of brain diseases. This is, as we demonstrate here, particular relevant for AD research.

## Acknowledgments

Work in the De Strooper laboratory was supported by the Fonds voor Wetenschappelijk Onderzoek (FWO), KU Leuven, VIB, UK-DRI (Medical Research council, ARUK and Alzheimer Society), a Methusalem grant from KU Leuven and the Flemish Government, Vlaams Initiatief voor Netwerken voor Dementie Onderzoek (VIND, Strategic Basic Research Grant 135043), the “Geneeskundige Stichting Koningin Elisabeth”, Opening the Future campaign of the Leuven Universitair Fonds (LUF), the Belgian Alzheimer Research Foundation (SAO-FRA) and the Alzheimer’s Association USA. Bart De Strooper is holder of the Bax-Vanluffelen Chair for Alzheimer’s Disease. Cell sorting was performed at the KU Leuven FACS core facility, and sequencing was carried out by the VIB Nucleomics Core. Renzo Mancuso is recipient of a postdoctoral fellowship from the Alzheimer’s Association, USA.

## Methods (online)

### *In vitro* generation of microglia from ESCs

In vitro microglia differentiation from embryonic stem cells was based on previously described protocols.^13,37^ On days 17, 21, 25, 28 and 32 of the protocol, non-adherent cells were harvested and selected using CD14-labelled magnetic beads (Miltenyi) following manufacturer specifications. Briefly, cells were collected and centrifuged for 5 min at 300g. Then, cells were incubated for 15 min at 4°C in 80ul MACS buffer (AUTOMACS + 5% MACS serum, Miltenyi) with 20ul of CD14-beads (Miltenyi), and passed through a LS column (QuadroMACS, Miltenyi). The CD14+ fraction was collected and centrifuged for 5min at 300g. Monocytes were then differentiated into microglia-like cells using microglia differentiation medium (TIC) (DMEM/F12, Glutamine (2mM), N-Acetyl Cysteine (5ug/mL), Insulin (1:2000), Apo-Transferrin (100ug/mL), Sodium Selenite (100ng/mL), Cholesterol (1.5ug/mL), Heparan Sulphate (1ug/mL)) supplemented with 50 ng/ml IL34, 50ng/mL M-CSF, 10ng/ml CX3CL1 and 25ng/mL TGF-β, resembling conditions described by Abud et al. (2017).^35^ The medium was changed every other day.

### Mice

*Rag2*^−/−^ *IL2rγ*^−/−^ *hCSF1*^*KI*^ mice^14^ were purchased from Jacksons Labs (strain 017708), and bred and maintained in local facilities. Mice were housed in groups of 2-5, under a 14 h light/10 h dark cycle at 21°C, with food and water *ad libitum*. All experiments were conducted according to protocols approved by the local Ethical Committee of Laboratory Animals of the KU Leuven (government licence LA1210591, ECD project number P177/2017) following local and EU guidelines.

### Endogenous mouse microglia depletion

The CSF1R inhibitor BLZ945 was dissolved in 20% (2-hydroxypropyl)-β-cyclodextrin (Sigma-Aldrich). Newborns were injected (i.p.) 24 and 48h prior to human cell transplantation at a dose of 200 mg/kg bodyweight.

### Transplantation of human microglia into the mouse brain

Grafting of human PSC-derived microglia was performed as previously described.^38^ Briefly, human microglia were dissociated and resuspended at a concentration of 100,000 cells/μl in PBS. At P4, mice were anaesthetised by hypothermia and bilaterally injected with 1μl of cell suspension at coordinates from Bregma: anteroposterior, −1mm; lateral, ±1mm. After the injections, mice were allowed to recover on a heating pad at 37 degrees, and then transferred back to their cage.

### Isolation of human primary microglia

Human primary microglia were isolated from brain tissue samples resected from the temporal cortex during neurosurgery. All samples represented lateral temporal neocortex and were obtained from patients who underwent amygdalohippocampectomy for medial temporal lobe seizures. The mesial temporal specimens were sent to pathology and thus not available for study purposes. Samples were collected at the time of surgery and immediately transferred to the lab for tissue processing, with post sampling intervals of 5-10 min. All procedures were conducted to protocols approved by the local Ethical Committee (protocol number S61186).

### Preparation and intracerebral injection of oligomeric amyloid

Oligomeric Aβ 1-42 (oAβ, 5μl 10μM) or scrambled peptide (Scr, 5μl 10μM) were prepared as previously described by Kuperstein et al.^39^ Briefly, recombinant amyloid beta 1-42 peptide (rPeptide; #A-1163-1) or scrambled amyloid beta 1-42 (rPeptide; #A-1004-1) thawed at room temperature 30 minutes before preparation. Then the peptides were solubilized in 99% hexafluoroisopropanol (HFIP) (Sigma-Aldrich; #105228) at 1 mg ml^−1^concentration. The HFIP was evaporated using a stream of nitrogen gas, the resulting peptide pellet is resolved in dimethylsulfoxide (DMSO; Sigma-Aldrich; #D4540), at final concertation of 1 mg ml^−1^. Next, the buffer exchange of DMSO with Tris-EDTA (50 mM Tris and 1 mM EDTA, pH 7.5) was performed using 5-ml HiTrapTM desalting columns (GE Healthcare; #17-408-01). The eluted peptide concentration was determined by using Bradford reagent (Bio-Rad; #5000006) according to the manufacturer’s instructions. The eluted peptide was left to oligomerize at room temperature for two hours in Tris-EDTA buffer. The oAβ or scrambled peptide was further diluted to 10 μM in Tris-EDTA buffer and stored at −80°C until use. At 8-10 weeks of age, grafted mice were anesthetized with a ketamine/xylazine mixture (85 and 13 mg/kg), and 5μl of either oAβ (10μM) or scrambled peptide (10μM) were stereotactically injected in the left ventricle at the following coordinates from bregma: anteroposterior, −0.1 mm; mediolateral, +1 mm; dorsoventral, −3 mm.^24^ Mice were allowed to recover in a thermoregulated chamber and then transferred back to their original cage. Isolation of microglia was performed 6h after the intracerebral injection of oAβ.

### Isolation of human and mouse microglia from the mouse brain

Mice were terminally anesthetized with an overdose of sodium pentobarbital and transcardially perfused with heparinized PBS. Brains were harvested in PBS 2% FCS 2mM EDTA (FACS buffer), mechanically triturated and enzymatically dissociated using the Neural Tissue Dissociation Kit (P) (Mylteni) following manufacturer specifications. Then, samples were passed through a cell strainer of 70μm mesh (BD2 Falcon) with FACS buffer, and centrifuged twice at 500g for 10 min at 4°C. Next, cells were resuspended in 35% Percoll (GE Healthcare) and centrifuged at 500g for 15 min at 4°C. The supernatant and myelin layers were discarded, and the cell pellet enriched in microglia was resuspended in FcR blocking solution (Mylteni) in cold FACS buffer, following manufacturer specifications. After a wash, primary antibody labeling was performed for 30 min at 4 °C, using the anti-CD11b (Mylteni) and anti-hCD45 (BD Bioscience), adding e780 (eBiocience) as a cell viability marker. Moreover, unstained cells and isotype-matched control samples were used to control for autofluorescence and/or non-specific binding of antibodies. Samples were run on a BD FACS Aria II Flow Cytometer and data were analysed using FlowJo and FCS express software. Human cells were sorted according to the expression of CD11b, hCD45, and GFP, whereas mouse cells only expressed CD11b but were negative for hCD45 and GFP (Supplementary Fig. 3). For each experimental condition, we pooled tissue from two mice.

### Histological analysis

Mice were terminally anesthetized with an overdose of sodium pentobarbital and transcardially perfused with heparinized PBS. Brains were harvested and cut in transverse serial sections (35μm thick) with a vibrating microtome (Leica). For each sample, 6 series of sections were sequentially collected in free-floating conditions and kept in cyoprotectant solution at −20°C. Sections were blocked with 5% normal serum in PBS PBS-0.2% Tween 20 for nonspecific binding. After rinses with PBS-0.1% Tween 20 (PBST), sections were incubated overnight at 4°C with anti-GFP, anti-Iba1 (Wako, 019-19741). After washes with PBST, sections were incubated with the appropriated biotinylated (Vector Labs) or Alexa 488- and 594-conjugated secondary antibodies (Invitrogen) for 1h at RT. When necessary, sections where incubated with Alexa 488-conjugated Streptavidin (Invitrogen) for 1h at RT. Finally, sections were counterstained with DAPI and mounted with Mowiol/DABCO (Sigma-Aldrich) mixture. Sections were visualized on a Nikon A1R Eclipse confocal system. Nearest neighbour distance (NND) analysis was performed in 20X microphotographs by using a script for Fiji (ImageJ) as previously described by Davis et al. (2017).^16^

### Single cell mRNA libraries preparation and sequencing

After microglial isolation, we performed single cell RNA sequencing by using 10X Genomics single cell gene expression profiling kit. cDNA libraries were produced following manufacturer instructions. cDNA libraries were then sequenced in an Illumina HiSeq platform 4000 with the sequencing specification recommended by 10X Genomics workflow. For each experimental condition, we pooled tissue from two mice.

### Human-mouse orthology

Human to mouse and mouse to human orthology tables were downloaded from Ensembl/Biomart (release 94)^9^. From these tables, only those genes were extracted that have a clean one-to-one ortholog in both directions. After filtering out genes that do not express in both our human and mouse microglia datasets, the table resulted in 10914 genes (Supplementary table 4).

### Analysis of single cell RNA sequencing datasets

#### Alignment

The raw BCL files were demultiplexed and aligned by Cellranger (version 2.1.1) against a human genome database (build hg38 build 84) and mouse database (mm10 build 84). Raw count matrices were imported in R (version 3.4.4) for data analysis.

#### Quality control of cells - step 1

For each dataset, to exclude poorly sequenced cells, damaged cells and dying cells, we filtered out cells with less than 100 genes detected; moreover, we excluded cells with more than 10% of reads aligning to mitochondrial genes. Genes detected in less than 3 cells were excluded from the count matrices. Data were analysed by principal component analysis (PCA) to identify any obvious batch effects. For the joint analysis of H9-derived microglia and primary microglia from surgical resections (Fig. 2a), the mean depth of sequencing was 102,000 reads/cell, while the mean number of genes detected per cell was 2072. For the analysis of mouse microglia (Fig. 3a), the mean depth of sequencing was 68,000 reads/cell, while the mean number of genes detected per cell was 1777. For the analysis of H9-derived microglia (Fig. 3f), the mean depth of sequencing was 96,000 reads/cell, while the mean number of genes detected per cell was 1964.

#### Quality control of cells - step 2

We analysed each dataset using the R package Seurat (version 2.3.4)^40^ for the mouse and H9-derived microglia datasets, and version 3.0^41^ for the joint analysis). We performed principal component analysis (PCA) on both the mouse and H9-derived microglia datasets, after data normalization and scaling and selection of the most variable genes, respectively 2000 and 1390. We selected the first principal components (PCs), 20 for mouse and 20 for H9-derived cells, based on a scree plot (i.e. a plot of the PC eigenvalues in decreasing order) as input for the downstream calculations. Clusters are identified using Seurat’s FindClusters function. Further non-linear dimensionality reduction for visualization is done using t-SNE. The standard workflow was followed also for the joint analysis, see Data integration and Joint clustering section.

In the joint dataset integrating H9-derived naive microglia and primary microglia from human cases, we identified 9 major cellular populations (Supplementary Fig. 5A). Clusters 0 to 4 expressed homeostatic microglia markers (Supplementary Fig. 5b) with cluster 2 overexpressing activated microglia markers (*APOE*), while cluster 5 expressed gene markers of brain resident macrophages (*MRC1*, *CD163*, *MAF*). Cluster 6 cells expressed high levels of oligodendrocytes markers (*MBP*), and were therefore removed from the count matrix. Cluster 7 expressed high levels of *CD14* and MHC type II genes, likely representing a population of monocytes. Clusters 8,9 expressed high levels of genes like *PTPRC* and low levels of microglia markers, possibly reflecting a neutrophil population, and were also removed from the final analysis. Post-QC a total of 10512 microglia, brain resident macrophages and monocytes cells were retained for further analysis.

For the mouse microglia dataset, we identified 12 major cellular populations, most of them showing a tight distribution on the t-SNE plot (Supplementary Fig. 6a), with two main clusters (6 and 7) clearly separating, as well as four other very small clusters (9,10,11,12). Clusters 0 to 5 expressed high levels of homeostatic microglia markers, which were instead not expressed in all the separated clusters (Supplementary Fig. 6b). Cluster 8 expressed activated microglia and cytokines markers (Supplementary Fig. 6b). Based on a panel of marker genes (Supplementary Fig. 6c), we could identify enrichment for markers of different cell types other than microglia in the six separated clusters. Clusters 6 and 7 showed high expression levels respectively of gene markers of neutrophils (*Ccrl2*) and monocytes (*Ccr2*). Clusters 9,10 and 12, all composed by very small number of cells, expressed gene signatures of other brain cells (astrocytes (*Clu*), neurons (*Npy*), oligodendrocytes (*Mbp*)). Cluster 11 was enriched in markers of cycling cells (*Top2a*). Overall, 89% of cells (13342/15036) in our post-QC dataset were microglia, and only these cells were retained for further analysis. The final analysis was performed on oAβ and scrambled peptide-treated cells (Fig. 3a), consisting of a final dataset of 9942 cells.

For the H9-derived microglia dataset, we identified 8 major cellular populations, distributed in two main groups of cells on the t-SNE plot (Supplementary Fig. 6d), both showing a treatment-associated distribution of cells (Supplementary Fig. 6e). Clusters 0,2,3,5 expressed homeostatic microglia markers (Supplementary Fig. 6f), while clusters 1,4 (26% of cells) expressed gene markers of brain resident macrophages (*MRC1*, *CD163*). Cluster 7 expressed low level of macrophage markers and some activation markers (*CD74*), while cluster 6 was enriched in markers of cycling cells (*MKI67*). Cluster 8 counted few cells, was very different from all the others and had no clear markers, probably reflecting a small population of doublets. Overall, 72% of cells (6444/8998) in our post-QC dataset were microglia, and only these cells were retained for further analysis. The final analysis was performed on oAβ and scrambled peptide-treated cells (Fig. 3f), consisting of a final dataset of 4880 cells.

#### Independent clustering of mouse and H9-derived microglia

Cells passing QC were analysed using functions provided with the Seurat package, version 2.3.4. Data was log normalized and we regressed out the variable of read count. Next, we identified the genes with highest variability and performed PCA on such gene set. We identified the most informative principal components based on a scree plot and we used these to perform cell clustering. Identification of differential expressed genes was performed using the Wilcox test implemented by Seurat’s FindMarker. t-SNE plots were prepared using Seurat’s t-SNE implementation. For the mouse microglia dataset, we considered 1020 highly variable genes for PCA and the first 15 PCs for clustering. The H9-derived human microglia dataset was analysed similarly as described above, by performing PCA on the 1886 most variable genes and by using the first 15 PCs to perform cluster analysis.

#### Data integration and Joint clustering

Cells passing QC were analysed using the Seurat package, version 3.0. The combined object (H9-derived naive microglia and primary microglia from patients) was split into a list, with each dataset as an element. Standard preprocessing (log-normalization) was performed individually for each of the two datasets, and variable features (nfeatures = 2000) that were identified based on a variance stabilizing transformation (selection.method = “vst”). Next, we identified anchors using the FindIntegrationAnchors function, giving the list of Seurat objects as input. We used all default parameters, including the dimensionality of the dataset (dims = 1:30). We passed these anchors to the IntegrateData function, in order to get an integrated (or ‘batch-corrected’) expression matrix for all cells, enabling them to be jointly analysed. We used the new integrated matrix for downstream analysis and visualization using the standard workflow.

#### Pseudotime analysis

To infer the pseudotime of microglia progression towards phenotypic change in response to oAβ challenge, we used the Monocle 2 package (version 2.6.4)^26,28^. We performed an unsupervised identification of cell trajectories and states, based on the top 200 marker genes identified with a differential expression analysis between oAβ treated cells and scramble-treated cells.

#### Differential Expression

Differential expression was performed using functions provided with the Seurat package; p values were calculated using the Wilcoxon rank-sum test. In Seurat’s function FindAllMarkers, no threshold for the min.pct parameter was applied, in order not to miss marker genes of rare cell populations. All the other parameters were set to default. Genes with adjusted p values (using a Bonferroni correction) < 0.05 were considered significantly differentially expressed. Differential expression was used to find cluster markers in all datasets. For Fig.3g, differential expression was performed with the FindMarkers function of Seurat comparing CRM and HM clusters, both in mouse and H9-derived human datasets, with no logFC or min.pct thresholds.

#### Z-scores of cell types signatures

For Fig. 2b, 3b and 3e, signatures were calculated using Seurat’s AddModuleScore function using a list of marker genes identified for each cell type or cell state: Homeostatic microglia signature (*CX3CR1*, *TMEM119*, *P2RY12*, *ELMO1*, *P2RY13*, *CSF1R*, *SFMBT2*, *IFNGR1*, *GPR34*, *SLC4A7*), Activated microglia signature (*APOE*, *SPP1*, *CSTB*, *LPL*, *TREM2*, *ITGAX*, *CST7*, *CLEC7A*, *GPNMB*, see Supplementary table 3), Cytokines response signature (*IL1B*, *CCL2*, *CCL3*, *CCL4*, *IL6*, *CXCL8*, *IL4I1*, *NFKB1*, *NFKB2*, *IL1A*), Macrophage signature (*MRC1*, *CD163*, *MAF*).

#### Pathway enrichment analysis

Pathway analysis was performed with GOrilla (Gene Ontology enRIchment anaLysis and visuaLizAtion tool)^42^, with single ranked list of genes as running mode. For both mouse and H9-derived microglia, genes were ranked by p-value adjusted taken from the Differential Expression analysis performed between CRM and HM clusters. The enriched ontology terms were then grouped by major functional categories, and the most significant terms (after multiple correction by FDR) in the H9-microglia dataset were compared to the same terms in the mouse host microglia dataset (Supplementary Fig. 10a). Each gene that was found significant in Differential Expression was then annotated with the functional categories it belongs to (Supplementary Fig. S10b, c), considering only the terms that were found significantly enriched in the GOrilla analysis.

### Data availability

Data from Karen-Shaul et al.^17^ is available from GEO (identifier GSE98969).

## Supplementary figures

**Figure S1.**
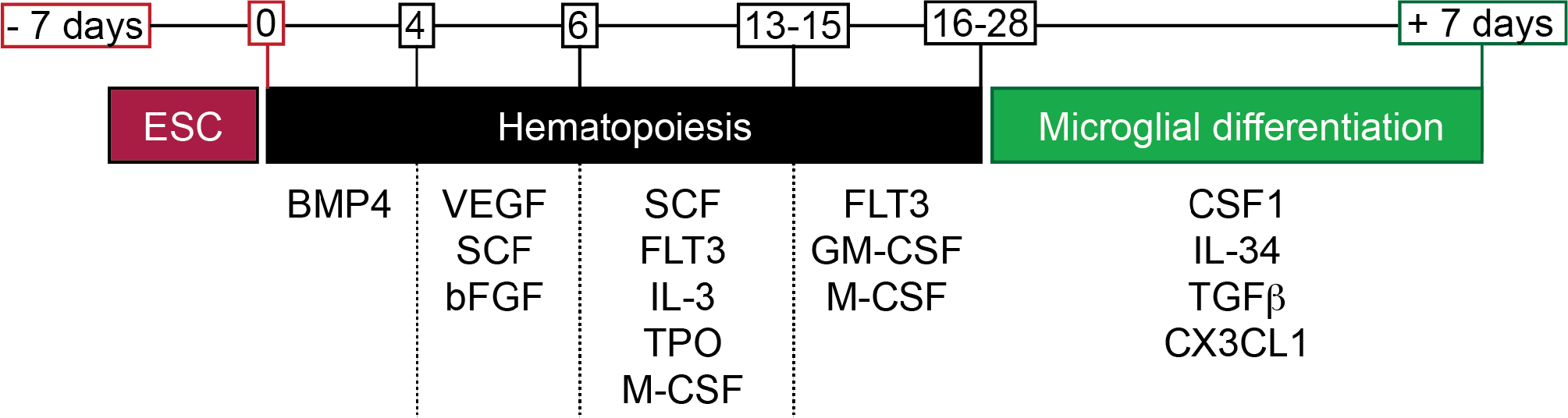
Schematic representation of the protocol followed to generate ESC-microglia like cells.

**Figure S2.**
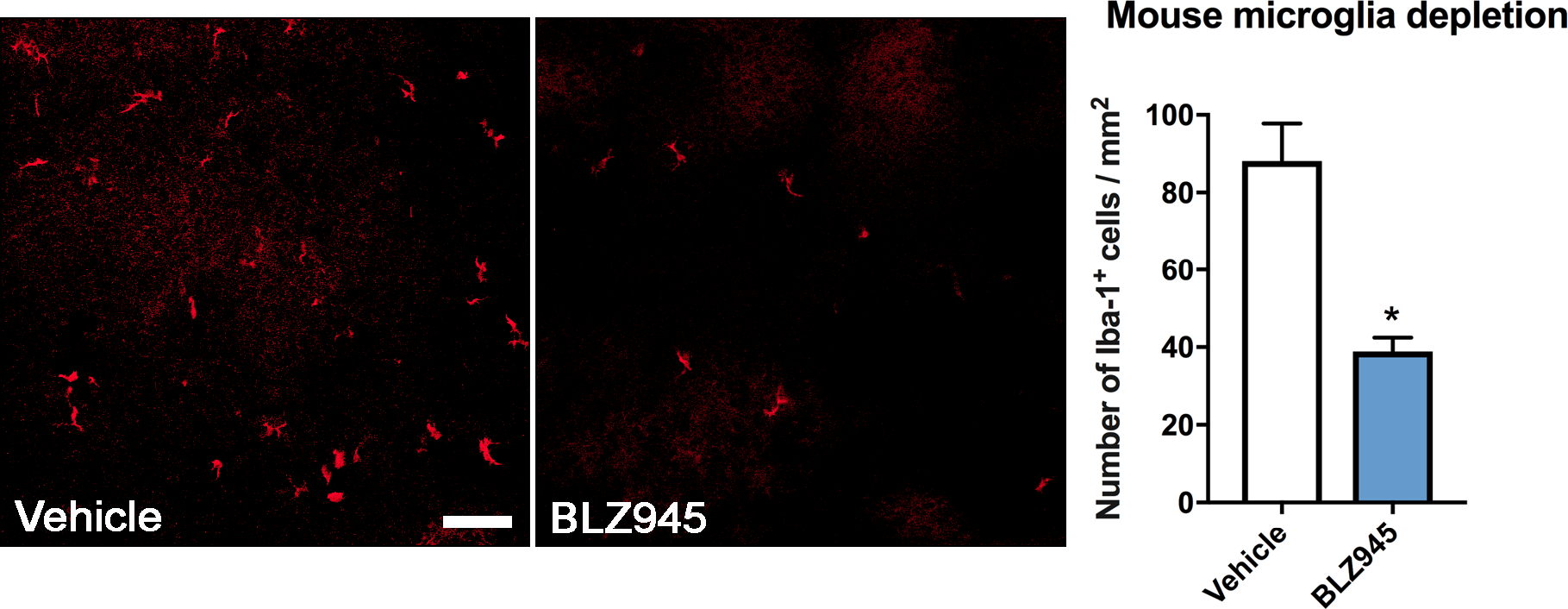
Endogenous mouse microglia depletion. CSF1R inhibitor BLZ945 was administered (200 mg/kg, i.p.) 24 and 48h prior to human microglia transplantation. Scale bar, 100μm. Graph shows mean±SEM, n=3 per group, two-tailed t-test *p<0.05.

**Figure S3.**
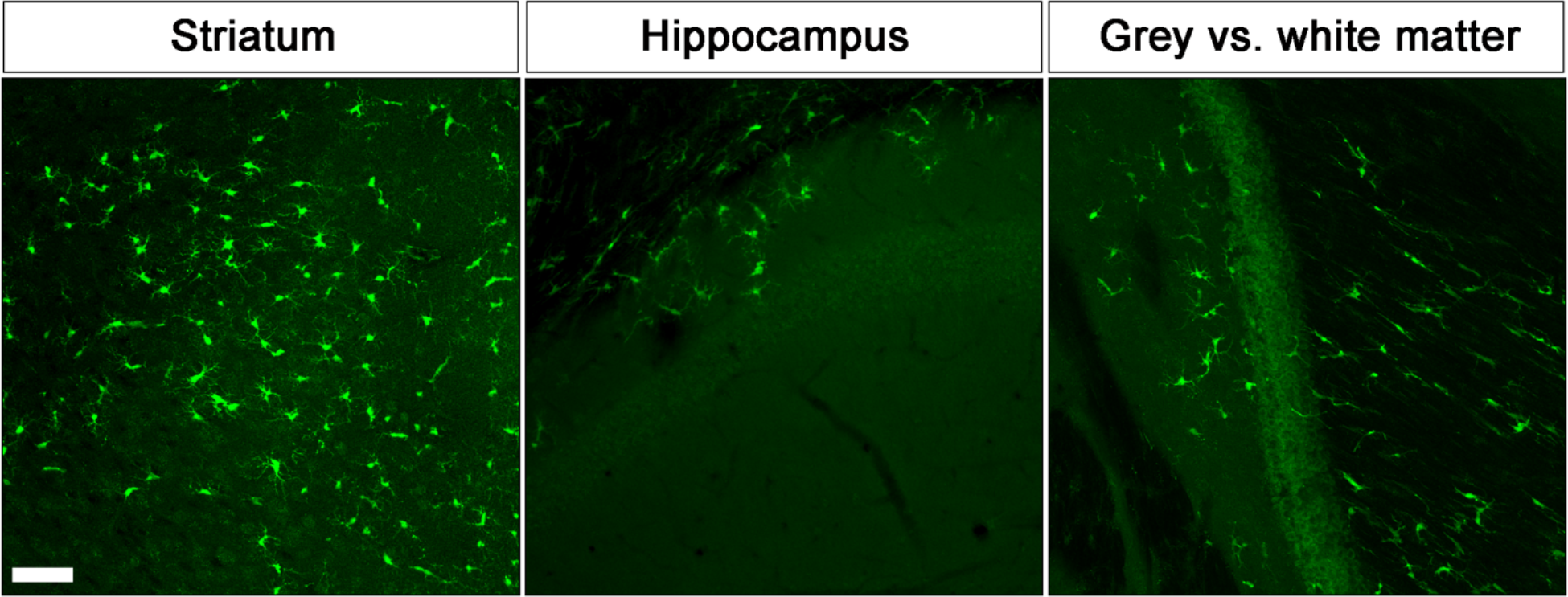
H9-microglia showed a widespread distribution across multiple areas of the brain. Representative images of (**a**) striatum (**b**) rostral hippocampus (**c**) fibre tracts. Note the different morphology displayed by the cells located in grey vs. white matter. Scale bar, 100μm.

**Figure S4.**
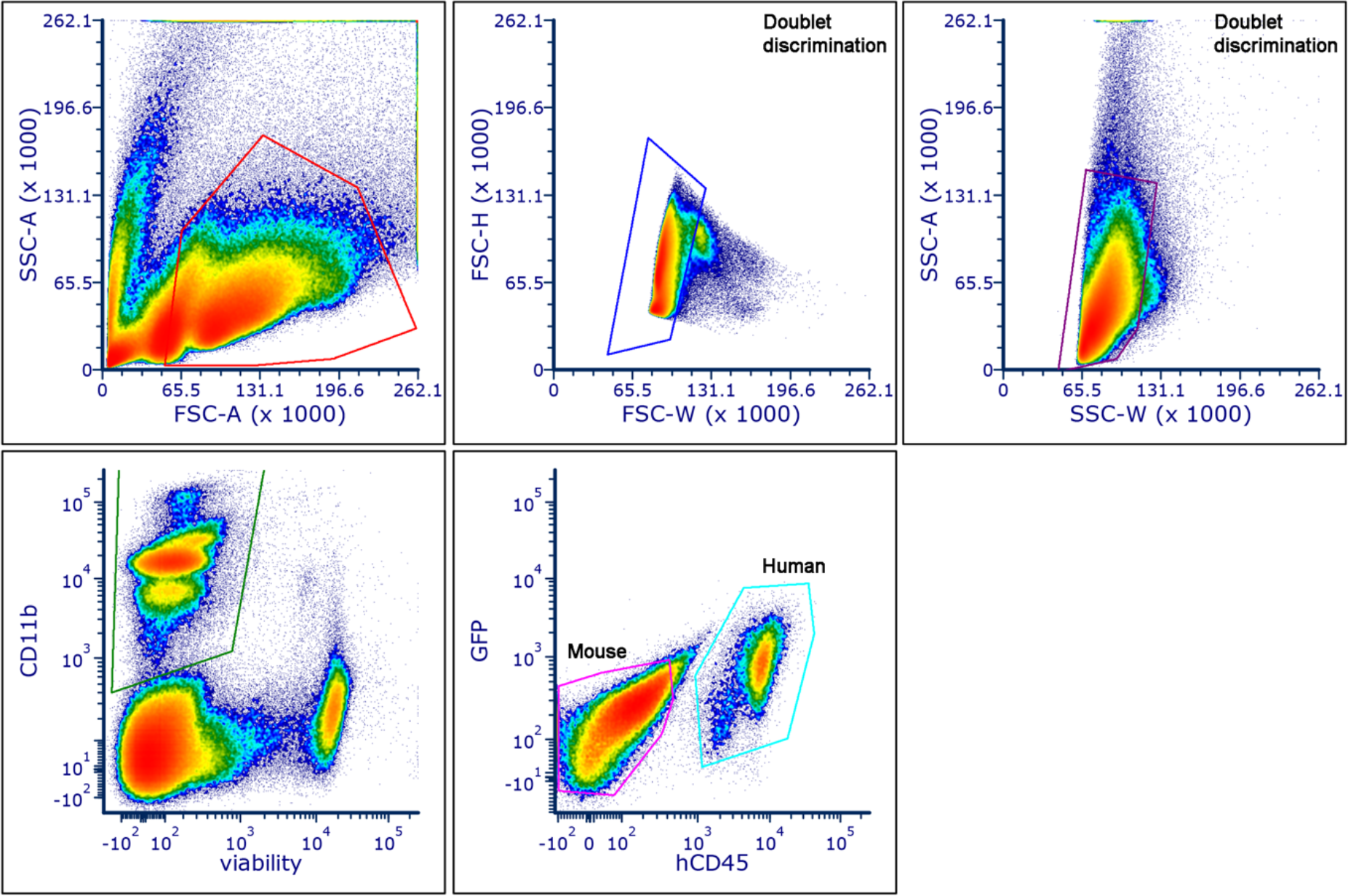
Gating strategy for the isolation of H9-microglia from the mouse brain. Human cells were sorted according to the expression of CD11b, hCD45, and GFP, whereas mouse cells only expressed CD11b but were negative for hCD45 and GFP.

**Figure S5.**
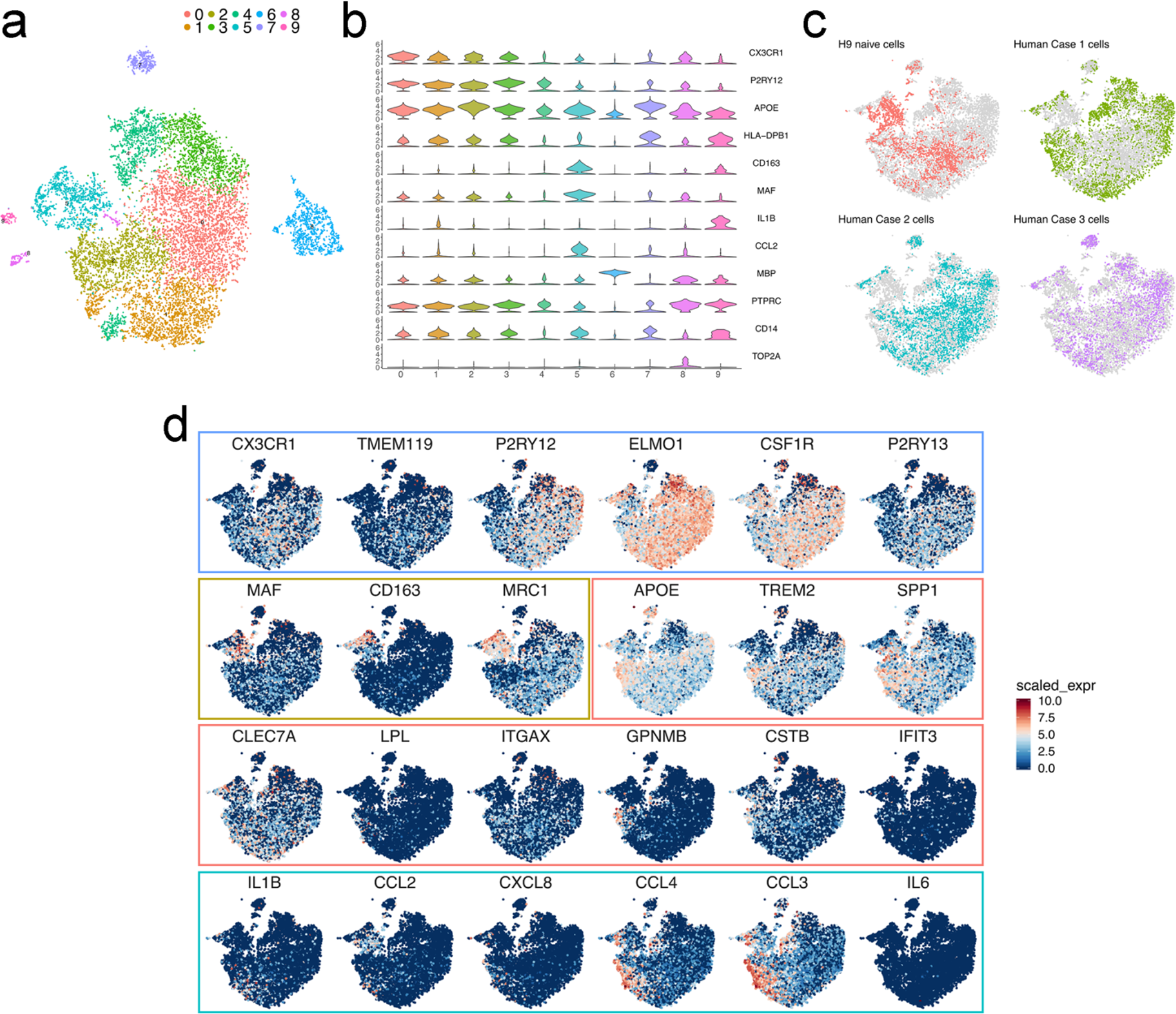
Preparation of the datasets for the analysis of naive H9-microglia and primary human microglia (see Fig. 2) (**a**) t-SNE plot of the cells passing quality control, coloured by clusters. (**b**) Violin plots of selected marker genes for homeostatic microglia (*CX3CR1*, *P2RY12*), ARM (*APOE*, *HLA-DRB1*), BMP (*CD163*, *MAF*), CRM (*IL1B*, *CCL2*), Oligodendrocytes (MBP), neutrophils (*PTPRC*), monocytes (*CD14*) and cycling cells (*TOP2A*). Analysis shown in **Fig. 2** was performed after removal of clusters 9 and 10 (neutrophils) and 12 (doublets). (**c**) Distribution of H9-microglia and each separate human case, superimposed onto the general cluster shown in Fig. 2a. (**d**) Selected genes defining the different transcriptomic scores shown in **Fig. 2b**: homeostatic microglia score in blue, macrophage score in brown, activated microglia score in red, and cytokines score in cyan. The colour scale represents the scaled expression for each gene.

**Figure S6.**
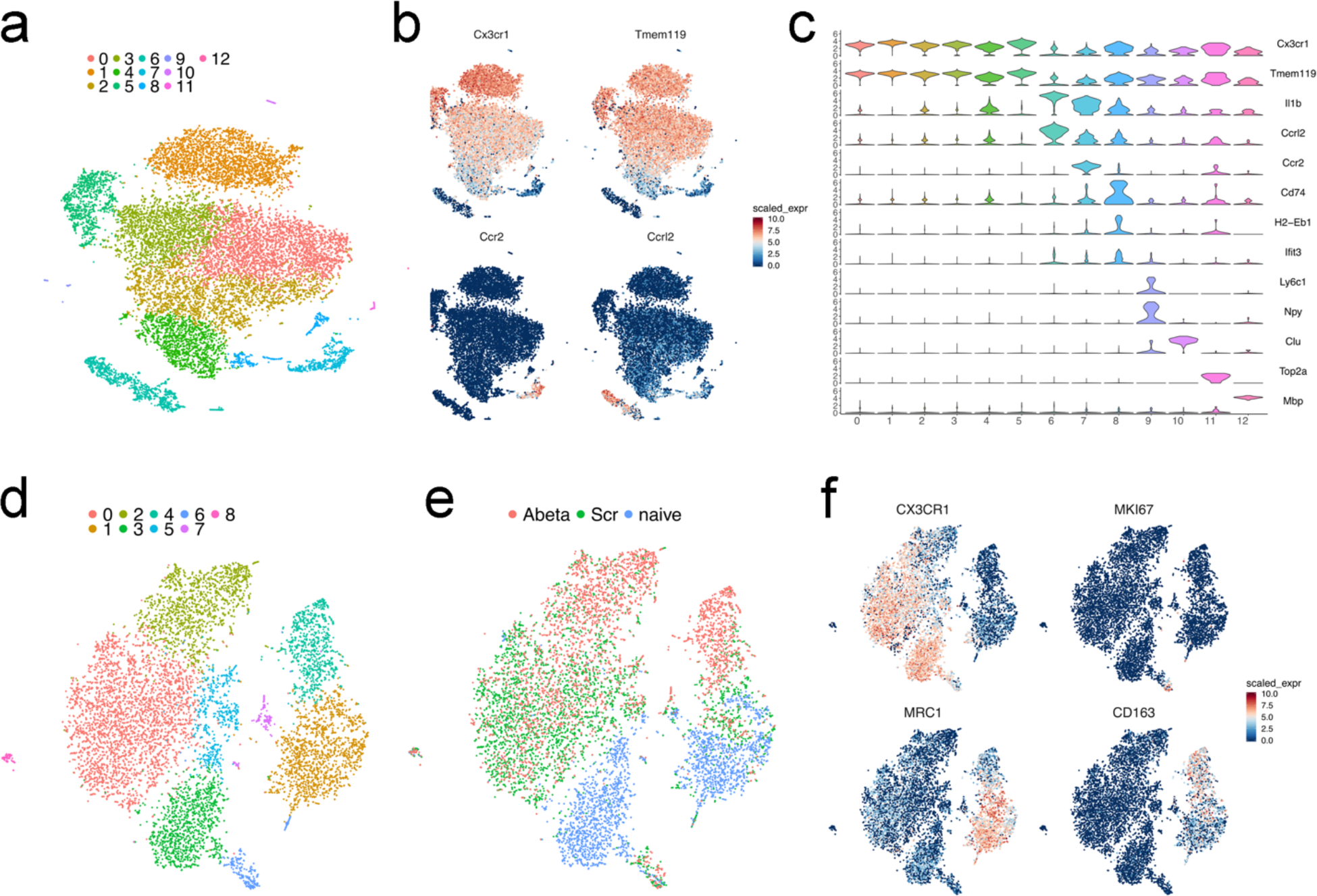
Preparation of the datasets for the analysis of (a-c) host mouse and (d-f) H9-microglia response to oAβ (see Fig. 3) (**a)** t-SNE plot of the cells passing quality control, coloured by clusters. (**b**) t-SNE plots as in **a**, coloured by the level of ln normalized expression of selected genes for microglia (*Cx3cr1*, *Tmem119*), monocytes (*Ccr2*) and neutrophils (*Ccrl2*). (**c**) Violin plots of selected marker genes for homeostatic microglia (*Cx3cr1*, *Tmem119*), CRM (*Il1b*), ARM (*Cd74*, *H2-Eb1*, *Ifit3*), neutrophils (*Ccrl2*), monocytes (*Ccr2*, *Ly6c1*), astrocytes (*Clu*), oligodendrocytes (*Mbp*), neurons (*Npy*), and cycling cells (*Top2a*). Analysis shown in **Fig. 2** was performed after removal of clusters 4 (neutrophils), 7 (monocytes), 10 (astrocytes), 12 (oligodendrocytes) and 9 (neurons). **(d)** t-SNE plot of the H9-microglia cells passing quality control, coloured by clusters. (**e**) t-SNE plot as in **a**, coloured by treatment (naïve; scrambled peptide, Scr; and oligomeric Aβ oAβ (**f**) t-SNE plot as in a, coloured by the level of ln normalized expression of selected genes for microglia (*CX3CR1*), cycling cells (*MKI67*) and brain resident macrophages (*MRC1*, *CD163*). Analysis shown in **Fig. 2** was performed after removal of clusters 1 and 4 (brain resident macrophages), 6 (cycling cells) and 8 (doublets).

**Figure S7.**
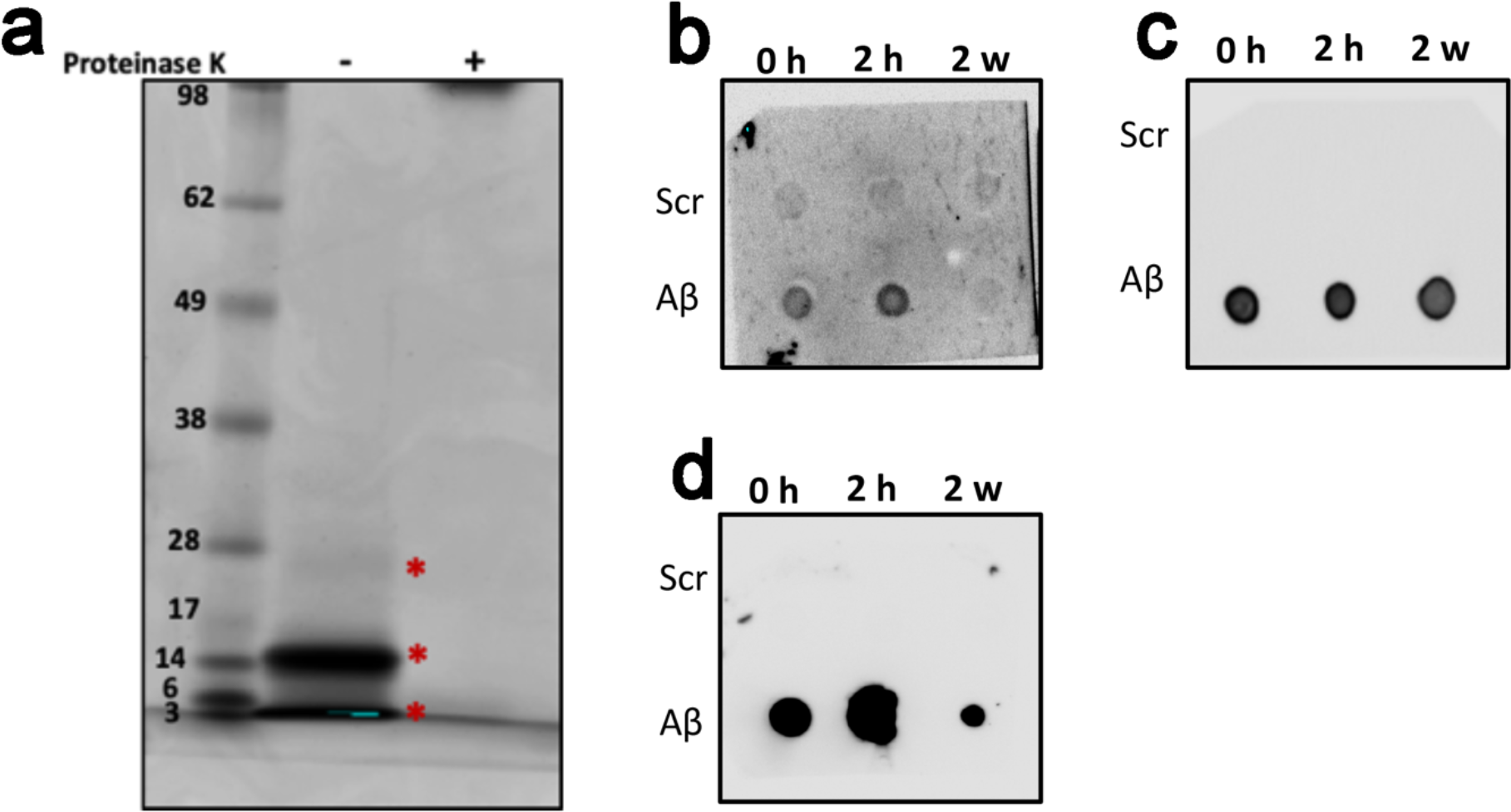
Characterization of oAβ preparation. Freshly eluted recombinant Aβ1-42 monomers follow a rapid aggregation course in Tris-EDTA buffer. (**a**) After 2 hours of incubation, Aβ1-42 monomers oligomerize and run as dimers and trimers (indicated with *) on SDS-PAGE/Coomassie staining, and they are proteinase-K sensitive. (**b**) Early Aβ1-42 oligomers form A11 and OC-positive aggregates. Two μl of either scrambled or amyloid beta 1-42 from different time points (0 hours, 2 hours and 2 weeks) of incubation was spotted on blots. These dot blots were probed with A11 antibody (Invitrogen; #AHB0052), which recognizes amino acid sequence-independent oligomers of proteins or peptides. A11 epitope is transient and is present only in the early oligomers (2 hours), in contrast to the mature fibers (2 weeks) formed after 2 weeks of incubation. No fibrillary material is detected after 2 hours of incubation. (**c**) OC antibody (Millipore; #AB2286) recognize epitopes common to monomers, amyloid oligomers, and fibrils. (**d**) 4G8 antibody (Eurogentec; #SIG-39220) detects N-terminal of the amyloid aggregates (epitope between amino acids 17-24).

**Figure S8.**
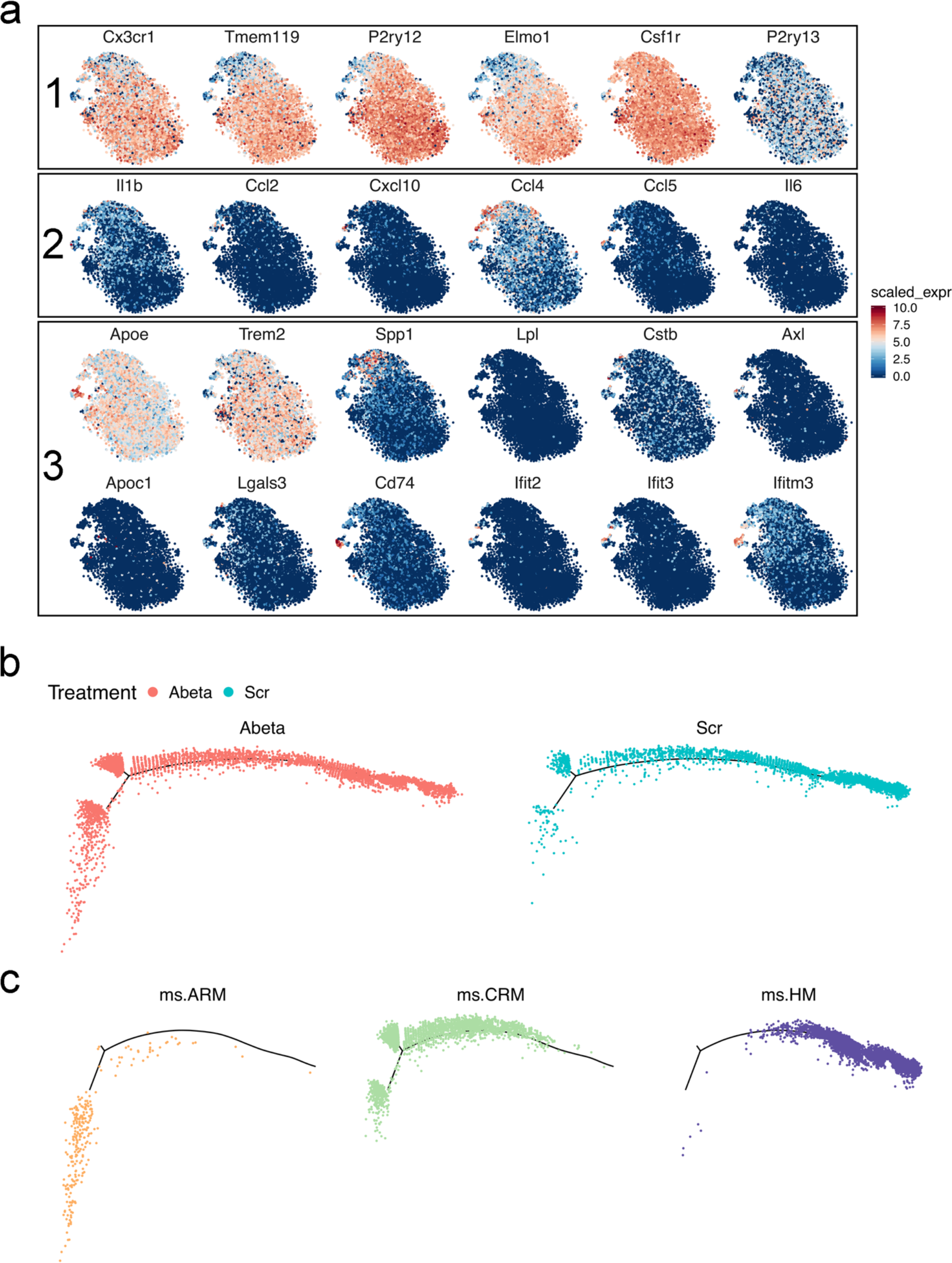
Expanded clustering and trajectory inference of the analysis of the response of mouse host microglia upon oAβ. (**a**) Selected genes defining the different transcriptomic scores shown in **Fig. 3b**: homeostatic score (1), cytokine score (2) activated score (3). (**b and c**) Phenotypic trajectory inferred by Monocle 2 as shown in **Fig. 3c**, coloured by (**b**) treatment and (**c**) clusters from **Fig. 3a**.

**Figure S9.**
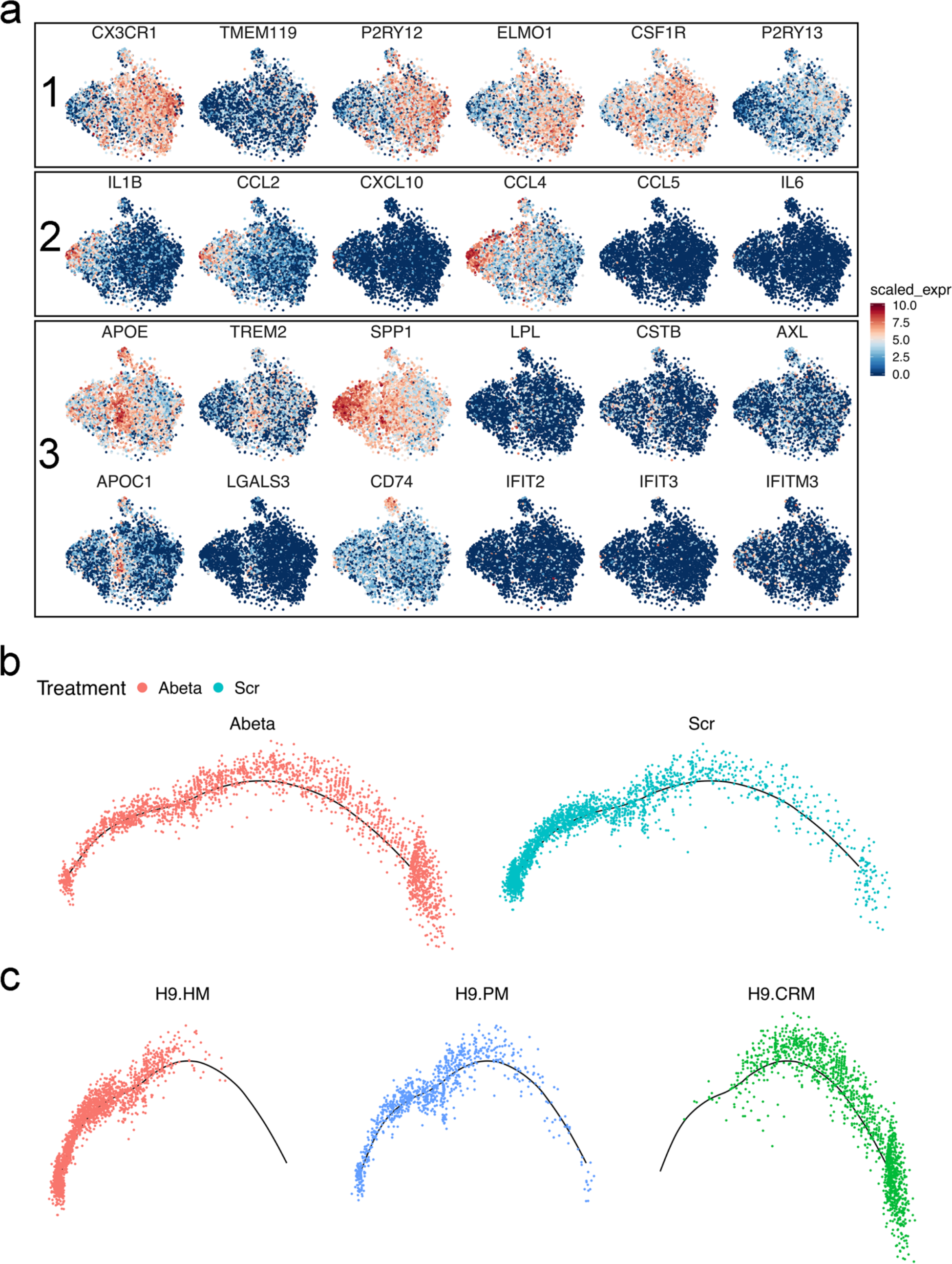
Expanded clustering and trajectory inference of the analysis of the response of H9-microglia upon oAβ. (**a**) Selected genes defining the different transcriptomic scores shown in **Fig. 3e**: homeostatic score (1), cytokine score (2) activated score (3). (**b and c**) Phenotypic trajectory inferred by Monocle 2 as shown in **Fig. 3d**, coloured by (**b**) treatment and (**c**) clusters from **Fig. 3f**.

**Figure S10.**
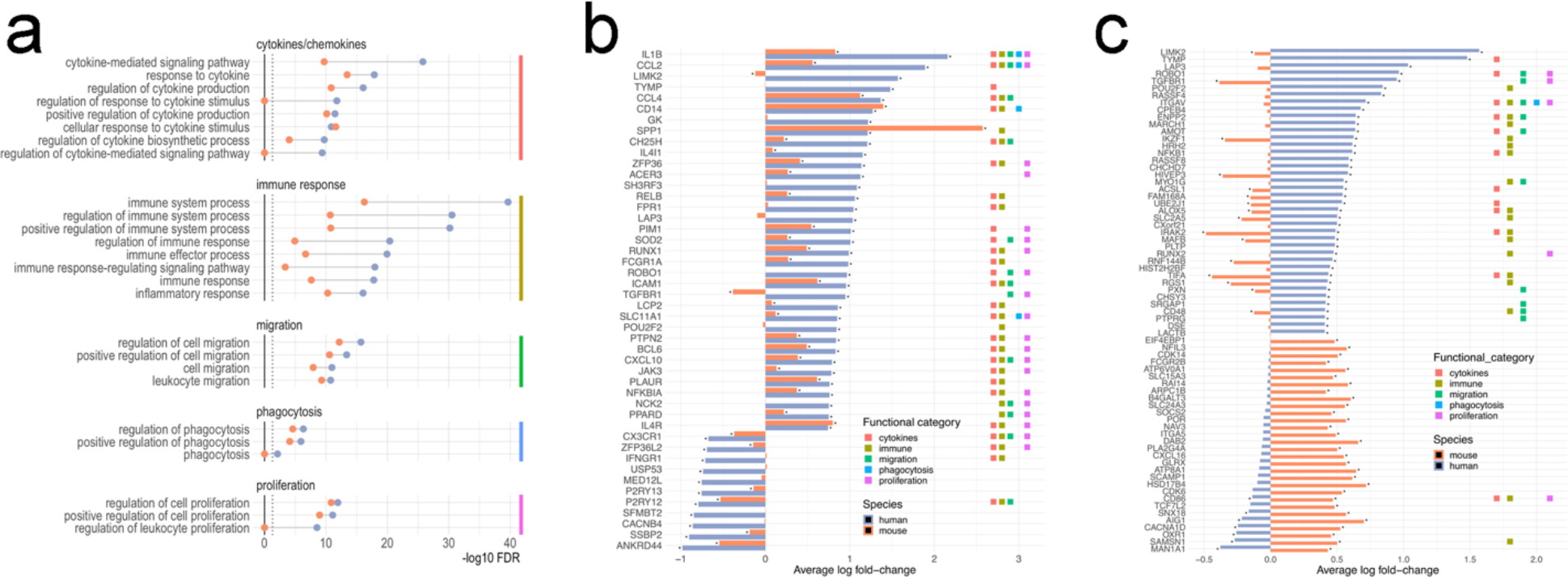
Differential responses of human and host mouse microglia to oligomeric Aβ. (**a**) Pathway enrichment analysis shows that the differentially expressed genes are involved in immune and inflammatory processes. (**b)** Top differentially expressed genes in H9-microglia upon Aβ challenge relative to scrambled peptide, and expression of their mouse orthologues by endogenous mouse cells. Coloured marks indicate the functional category as shown in **b**. (**c**) Differentially expressed genes that show opposite behaviour in H9- and mouse host (Rag2^−/−^) microglia. Coloured marks indicate the functional category as shown in **b**.

